## Supplementary Figures for "TRPA1 activation in non-sensory supporting cells contributes to regulation of cochlear sensitivity after acoustic trauma"

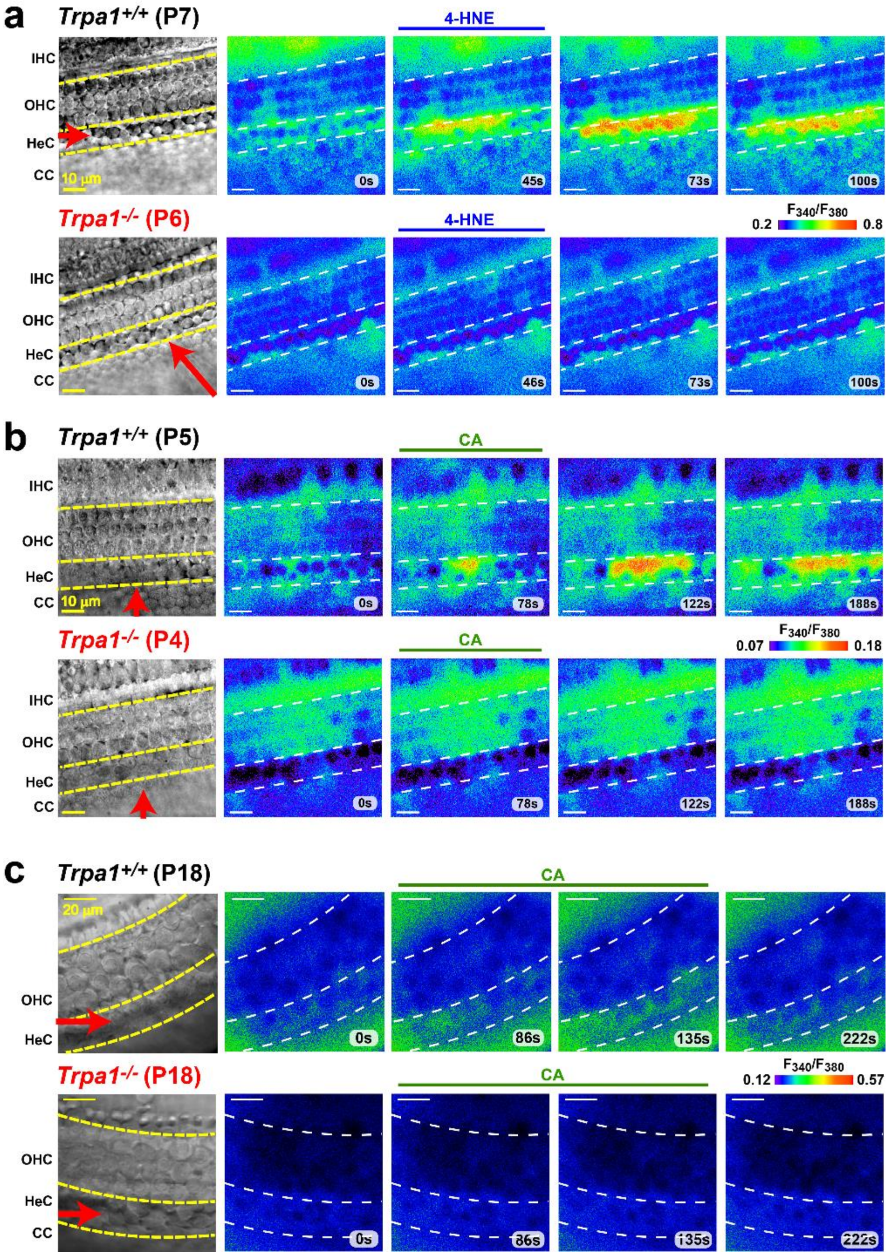


**Supplementary Figure 1.** ***Long-lasting $Ca^{2+}$ responses in Hensen's cells evoked by the application of TRPA1 agonists***

**(a,b,c)** Ratiometric ($F_{340}/F_{380}$) fura-2 images before normalization to the background $F_{340}/F_{380}$ values for the time-lapse frames presented in Fig.2a,d,g. Layout of the panels is identical to

Fig.2a,d,g except that the pseudocolor scale represents the $F_{340}/F_{380}$ ratio. This ratio is proportional to the absolute concentration of free cytosolic $Ca^{2+}$.

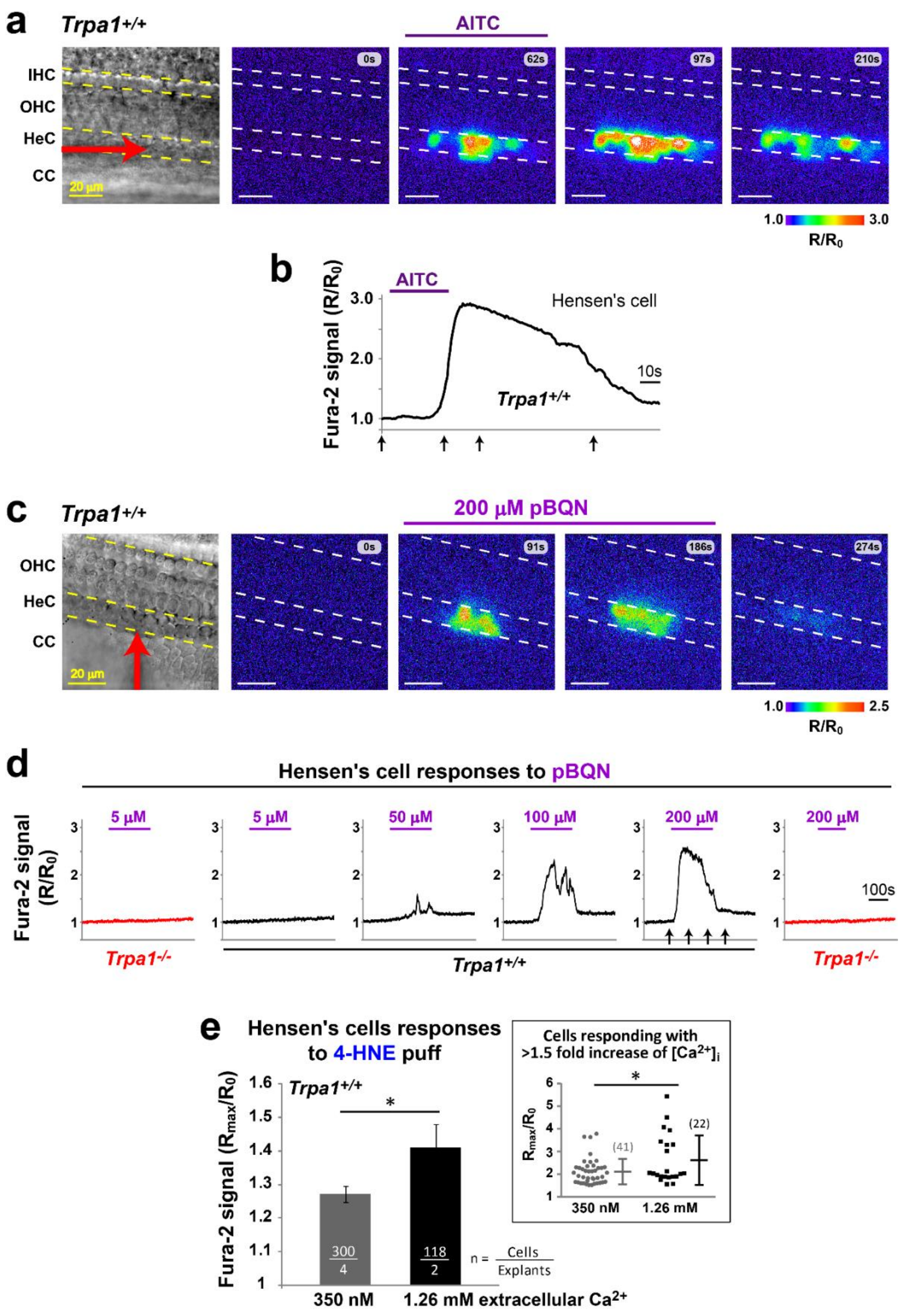


**Supplementary Figure 2.** ***TRPA1-mediated $Ca^{2+}$ responses in Hensen's cells to puff applications of AITC and pBQN***

**(a)** Time-lapse imaging of $Ca^{2+}$ responses to the application of mustard oil (AITC, 100 μM) in a cochlear explant from a wild-type P1 mouse. The reference bright field image indicating the position of the puff pipette (red arrow) is on the left, while other frames show the $R = F_{340}/F_{380}$ ratio of fura-2 signals in pseudocolor scale normalized to the pre-stimulus baseline ($R_0$). **(b)** Representative $Ca^{2+}$ response in one of the Hensen's cells shown in **a**. Arrows at x-axis indicate the time points when the imaging frames in **a** were taken. **(c)** $Ca^{2+}$ responses ($R/R_0$) to the local application of 200 μM *para*-benzoquinone (pBQN) in a cochlear explant from a P6 wild-type mouse. A reference bright field image of the same field of view is shown on the left. The yellow arrow indicates the position and direction of the puff stimulation. **(d)** Representative $Ca^{2+}$ responses to varying concentrations of pBQN in wild-type (black) and *Trpa1*$^{-/-}$ (red) Hensen's cells. Arrows in the x-axis of the $Ca^{2+}$ response to 200 μM benzoquinone indicate the time points where the frames in **c** were taken. **(e)** Normalized amplitude ($R_{max}/R_0$) of $Ca^{2+}$ responses to 200 μM 4-HNE puff application in wild-type Hensen's cells in the presence of either 350 nM or 1.26 mM extracellular $Ca^{2+}$. The data are shown as Mean±SE, and the asterisk indicates statistical significance ($P<0.05$, Student's *t* test). The inset shows the subset of these data from the cells responding with at least a 1.5 fold increase of $[Ca^{2+}]_i$, and the error bars indicate Mean±SD.

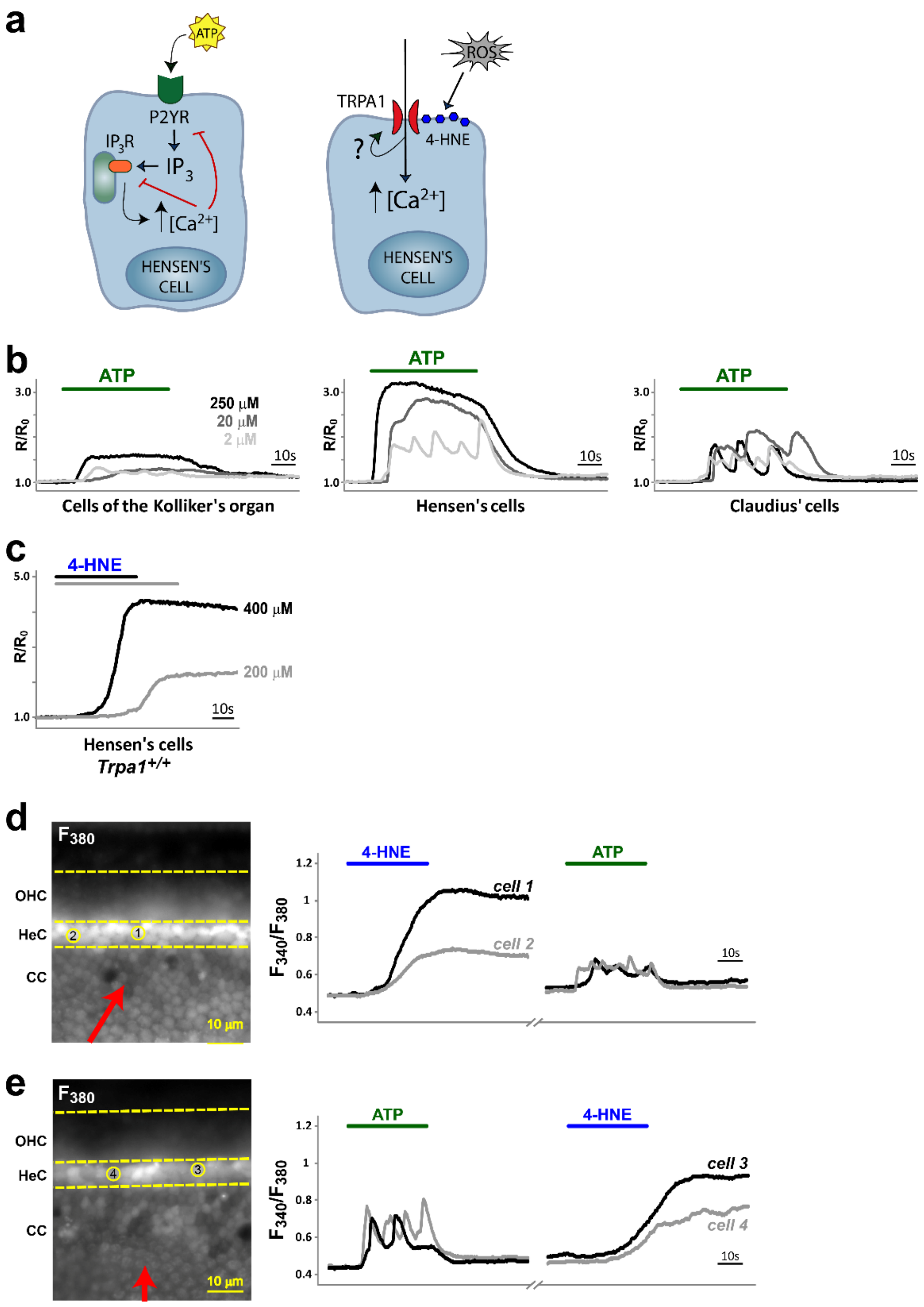


**Supplementary Figure 3.** ***Different kinetics of Ca$^{2+}$responses evoked by extracellular ATP and TRPA1 agonists in supporting cells of wild-type mice***

**(a)** Potential mechanisms of Ca$^{2+}$ responses to extracellular ATP (*left*) and TRPA1 agonists (*right*). *Left*, at low ATP concentrations, oscillating Ca$^{2+}$ responses have been associated with activation of G-protein-coupled ATP receptors (P2YR) and their subsequent desensitization, as well as with the $Ca^{2+}$-dependent feedback inhibition of $Ca^{2+}$ release from the intracellular stores[30]. *Right*, a TRPA1 channel may be 'locked' in an open state due to covalent modifications of TRPA1 by a reactive agonist (*e.g*. 4-HNE) or due to the potentiation of TRPA1 activation by intracellular $Ca^{2+}$[8, 9]. **(b)** Representative $Ca^{2+}$ responses to the application of 2, 20 and 250 µM ATP (darker traces indicate higher ATP concentrations) in Kolliker's organ (*left*), Hensen's cells (*middle*) and Claudius' cells (*right*). **(c)** Representative $Ca^{2+}$ responses to the puff application of 200 µM (gray) and 400 µM (black) of 4-HNE in Hensen's cells. Note that even the decreased response to 4-HNE at 200 µM does not show oscillations and continue long after the end of agonist application. **(d,e)** $Ca^{2+}$ responses in two Hensen's cells (black and grey) of a wild-type mouse evoked by the application of 4-HNE (200 µM) followed by ATP (2 µM) (**d**) or vice versa (**e**). Left panels show $F_{380}$ images of the analyzed cells (circled numbers) and positioning of the puff pipettes (red arrows). Breaks in the time axes on the right graphs represent the washout periods after the first stimulation. During this washout, the puff pipette was carefully replaced to deliver a new drug from the same position. Notice that the same cells exhibited oscillating short-lived $Ca^{2+}$ responses to ATP but long-lasting $Ca^{2+}$ responses to 4-HNE, regardless of the order in which these stimuli were delivered.

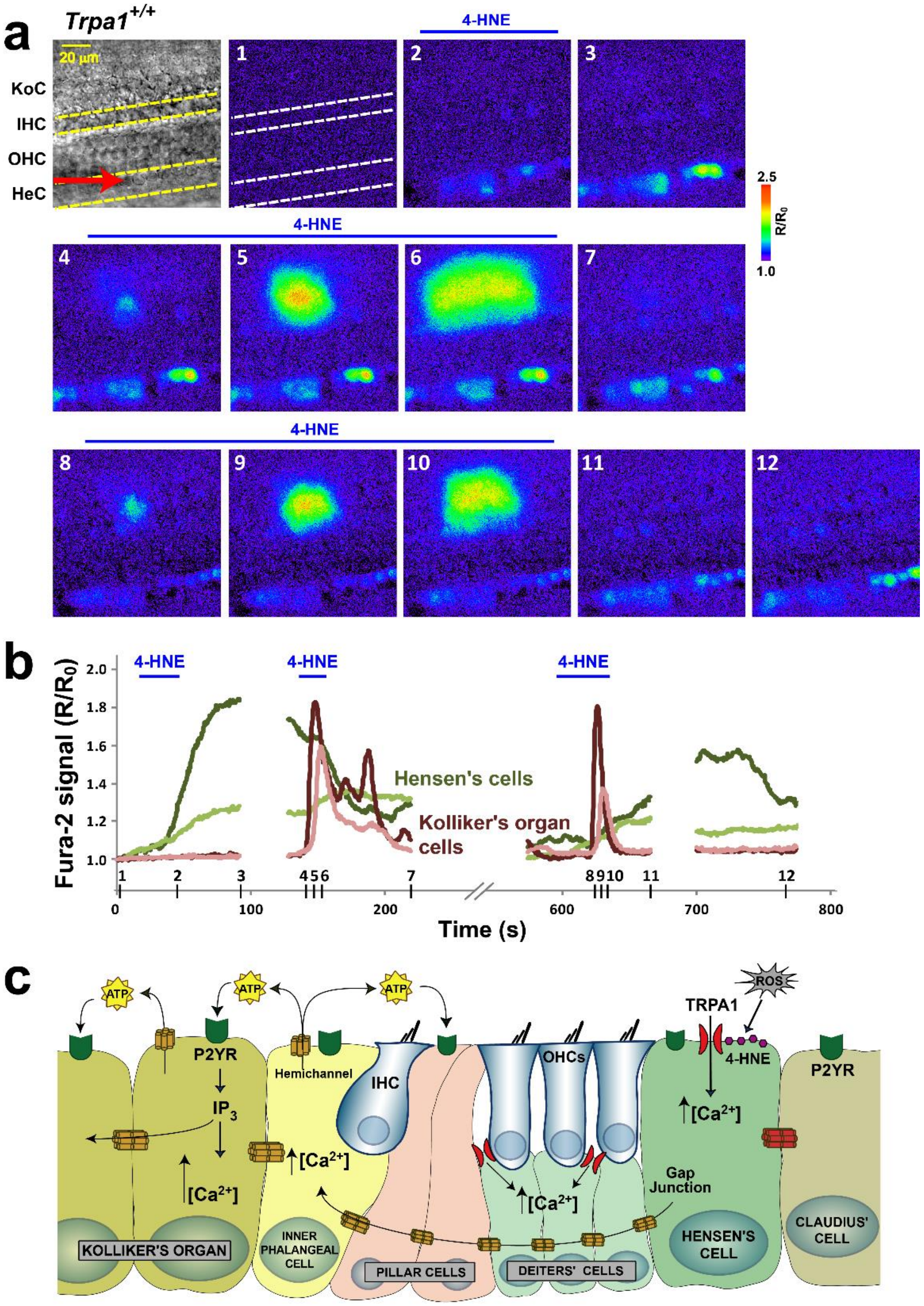


**Supplementary Figure 4.** ***TRPA1-initiated $Ca^{2+}$ responses propagate from Hensen's cells toward the Kolliker's organ at "hot spots"***

**(a)** Consecutive applications of 400 µM of 4-HNE to a Hensen's cell region evoke $Ca^{2+}$ responses propagating to the Kolliker's organ. First panel shows a reference bright field image with the puff pipette position (red arrow). Consecutive 4-HNE applications are indicated with blue horizontal bars above the images. Notice that the second and third stimulations of Hensen's cells led to the propagation of $Ca^{2+}$ responses to the same area of the Kolliker's organ. **(b)** Changes of $[Ca^{2+}]_i$ in two Hensen's cells (light and dark green) and two cells of the Kolliker's organ (light and dark red) from the experiment shown in **a**. The numbered ticks at the x-axis indicate the time points where the twelve frames in **a** were taken. **(c)** Potential mechanism of propagation of TRPA1-initiated signals. On the endolymphatic side of the epithelium, TRPA1 channels in Hensen's cells are the first to respond to endogenous byproducts of oxidative stress such as 4-HNE. Long-lasting $Ca^{2+}$ responses in Hensen's cells do not activate adjacent Claudius' cells but, instead, propagate across the organ of Corti and trigger $Ca^{2+}$ waves in the Kolliker's organ. These 'secondary' $Ca^{2+}$ waves in the Kolliker's organ depend on binding of extracellular ATP to P2Y receptors, thus resembling the $Ca^{2+}$ waves occurring after OHC damage. Propagation of $Ca^{2+}$ responses from Hensen's cells to the Kolliker's organ most likely involves the gap-junctional conductance through Deiters' and pillar cells. Rise of intracellular $Ca^{2+}$ can trigger ATP release to the extracellular space from cells of the Kolliker's organ. Extracellular ATP can bind P2Y receptors in various supporting cells, resulting in the downstream activation of TRPA1 channels on the perilymphatic side of the epithelium, thereby amplifying $Ca^{2+}$ responses (see also Fig.4). TRPA1 channels in Deiters' cells may sense byproducts of oxidative stress in the perilymph.

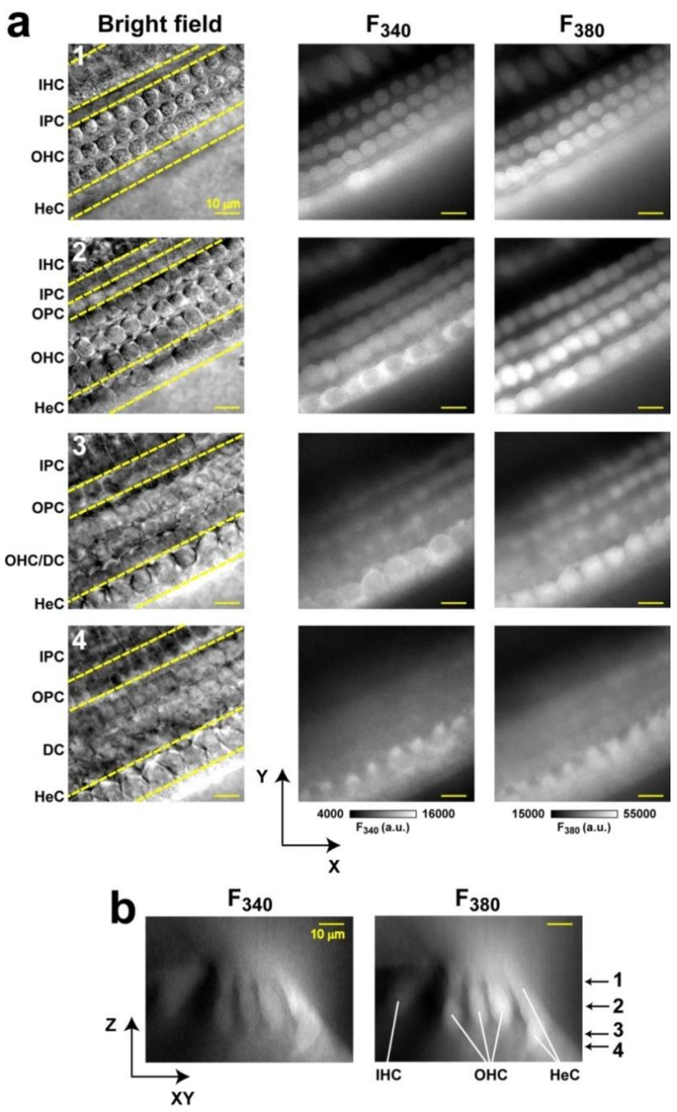


**Supplementary Figure 5.** ***Non-uniform loading of the cochlear epithelium with membrane-permeable fura-2***

**(a)** *Z*-stack of bright-field (*left*) and fura-2 signals in a cochlear explant obtained with the 340 (*middle*) and 380 (*right*) nm illumination. Reference bright-field images show the boundaries between different types of cells in the cochlear epithelium. **(b)** Orthogonal sections of fura-2 fluorescence reconstructed from the data shown in **a**. The arrows on the far-right side indicate the focal planes of the images shown in **a**. Notice the lack of fluorescence in both channels ($F_{340}$ and $F_{380}$) in the pillar cell area between the inner and outer hair cells and in Deiters' cells below OHCs.

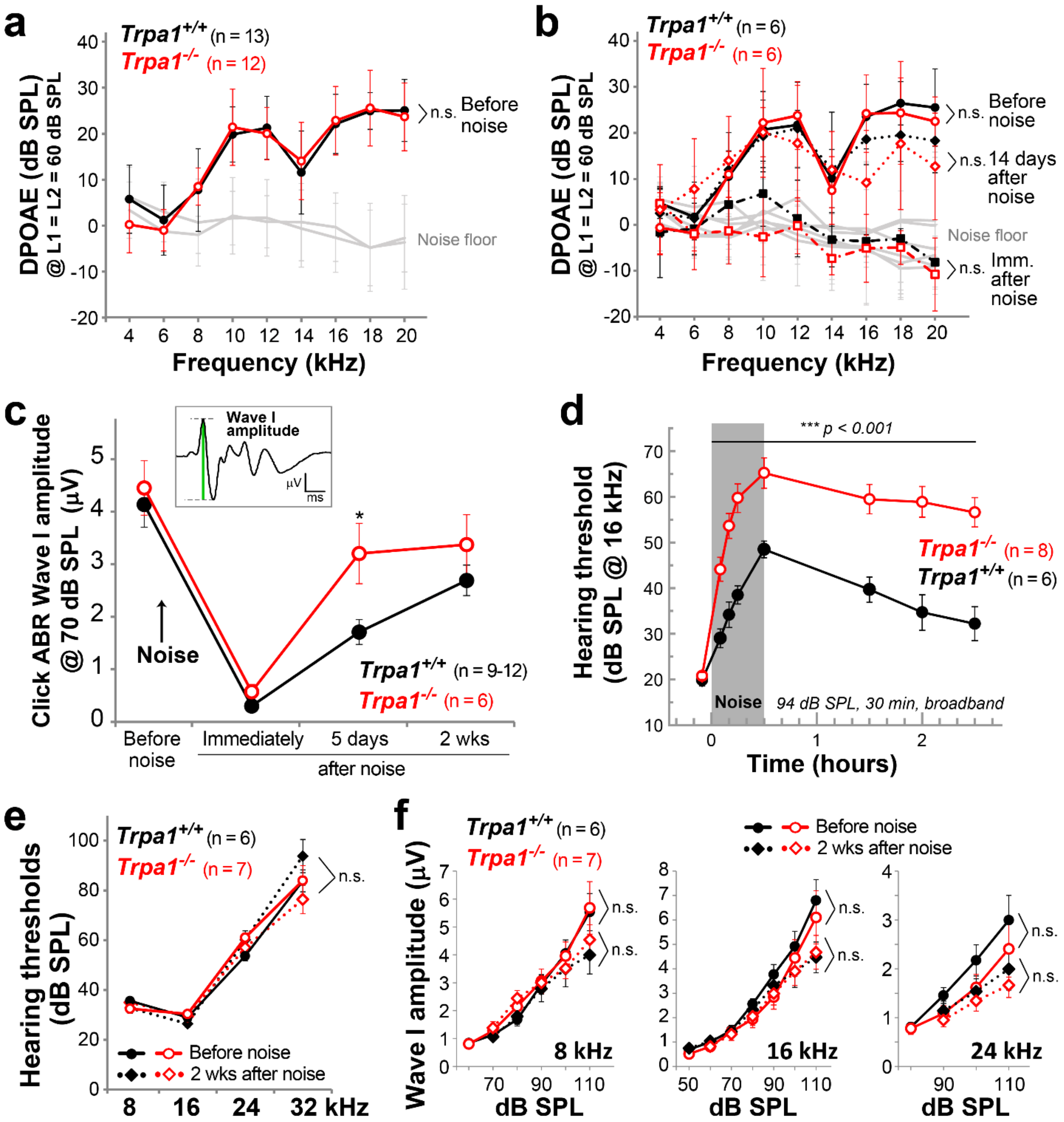


**Supplementary Figure 6.** ***Effects of moderate vs. mild noise exposures on hearing in mice lacking TRPA1 channels.***

**(a)** Baseline distortion product otoacoustic emissions (DPOAE) at different frequencies in wild-type (black, filled symbols) and *Trpa1*$^{-/-}$ mice (red, open circles) before noise exposure. **(b)** In a subset of experiments from panel (a), DPOAEs were measured before, immediately after (squares, dash-dot lines), or 14 days after (diamonds, dotted lines) exposure to moderate broadband noise (100 dB SPL, 30 min). **(c)** Amplitude of click-evoked ABR Wave I (as indicated in cartoon) in wild-type (filled symbols) and *Trpa1*$^{-/-}$ (open symbols) mice after moderate noise exposure (100 dB SPL for 30 min). **(d)** Hearing thresholds in wild-type (black, filled symbols) and *Trpa1*$^{-/-}$ (red, open symbols) mice determined with 16 kHz tone burst-evoked auditory brainstem responses (ABR) at several time points during and after exposure to moderate broadband noise (94 dB SPL, 30 min). **(e)** Thresholds of tone burst-evoked ABRs in wild-type (black, filled symbols) and *Trpa1*$^{-/-}$ (red, open symbols) mice before (circles, solid lines) and 2 weeks after (diamonds, dotted lines) exposure to mild broadband noise (85 dB SPL, 30 min). **(f)** Amplitudes of tone burst-evoked ABR Wave I in wild-type (filled symbols) and *Trpa1*$^{-/-}$ (open symbols) mice before (circles, solid lines) and 2 weeks after (diamonds, dotted lines) exposure to mild broadband noise (85 dB SPL for 30 min). Data are shown as Mean ± SD (**a**,**b**) or Mean ± SE (**c**,**d**,**e,f**). Asterisks indicate statistical significance (∗, $P<0.05$; ∗∗, $P<0.01$; ∗∗∗, $P<0.001$) at specific data points between genotypes (**c**,**d**) by Student's *t* test. Differences between grouped data (**a,b,e,f**) were assessed by two-way ANOVA and are shown to the right of traces; *n.s.* not significant.

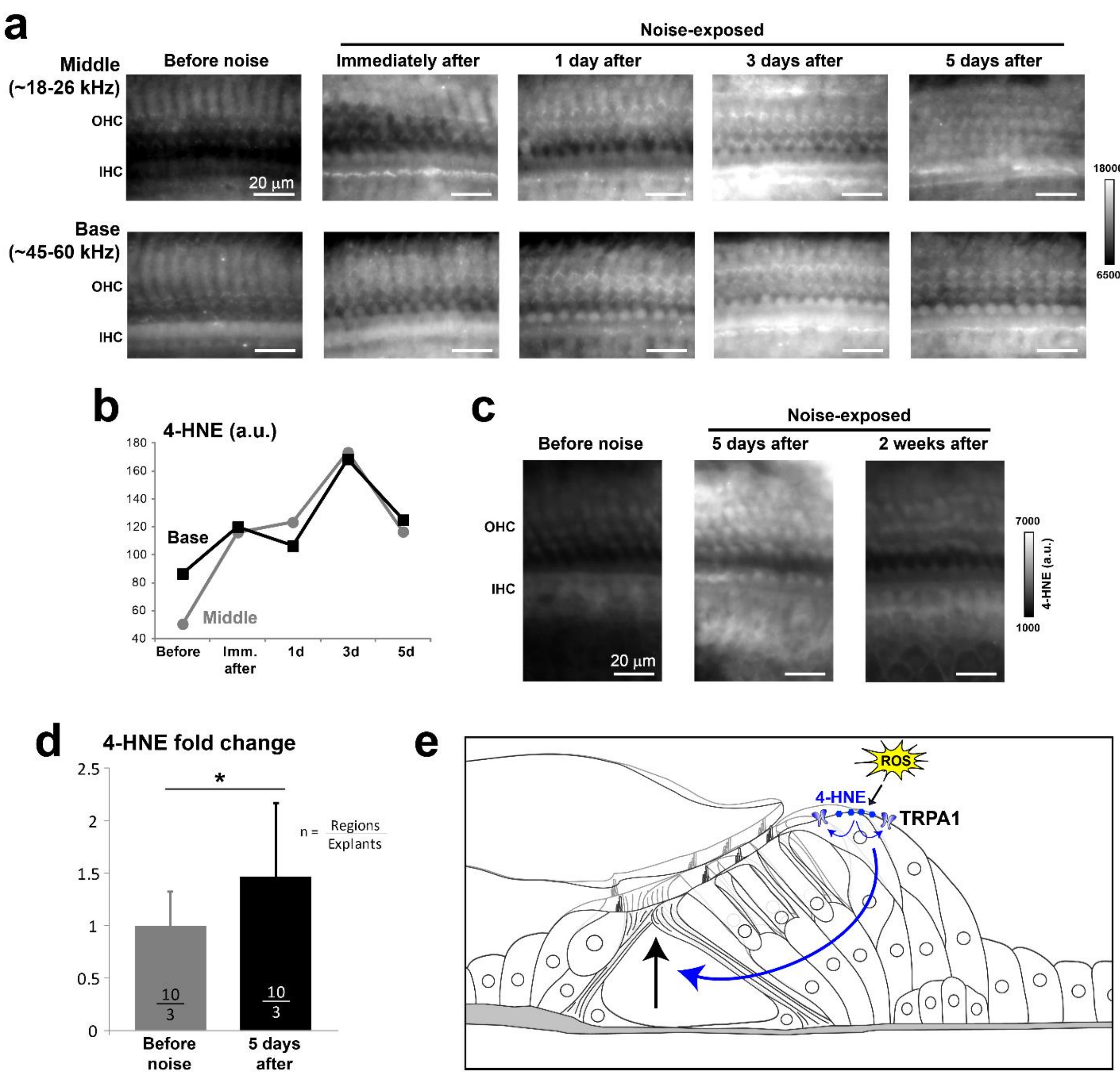


**Supplementary Figure 7.** ***Delayed increase of 4-HNE-modified proteins following noise exposure***

**(a)** Immunolabeling of 4-HNE-modified proteins in the organ of Corti from the middle (*top*) and basal (*bottom*) cochlear turns from 4-week old mice before and at several time points after a single exposure to wide-band noise at 100 dB SPL for 30 min. **(b)** Quantification of the 4-HNE fluorescence (a.u., arbitrary units) from the panels shown in **a**, indicating the progressive increase of oxidative stress byproducts throughout several days after noise exposure. **(c)** Immunolabeling of 4-HNE-modified proteins in the organ of Corti of 3-week old mice before (*left*), 5 days after (*middle*), and 2 weeks after (*right*) noise exposure, showing the eventual decrease of 4-HNE production by the second week after the acoustic trauma. Each image represents the average of a Z-stack covering the thickness of the organ of Corti. **(d)** Quantification of 4-HNE labeling before and 5 days after noise exposure in three independent series (two exposed to 100 dB SPL for 30 min and one to 110 dB SPL for 2 hours). Data are shown as Mean±SE, and the asterisk indicates statistical significance ($P<0.05$, Paired student's *t* test). **(e)** For several days after the noise exposure, the increase in reactive oxygen species (ROS) generates byproducts, such as 4-HNE, that can activate TRPA1 channels. 4-HNE causes robust and long-lasting $Ca^{2+}$ responses in the Hensen's cells. Then, Deiters' cells become activated either through propagation of the signal from Hensen's cells or by delayed activation by 4-HNE. Finally, the signal propagates to pillar cells and evokes combined changes in the shape of pillar and Deiters' cells, which presumably maintains a modified geometry of the organ of Corti and elevated hearing thresholds for days after the noise exposure.

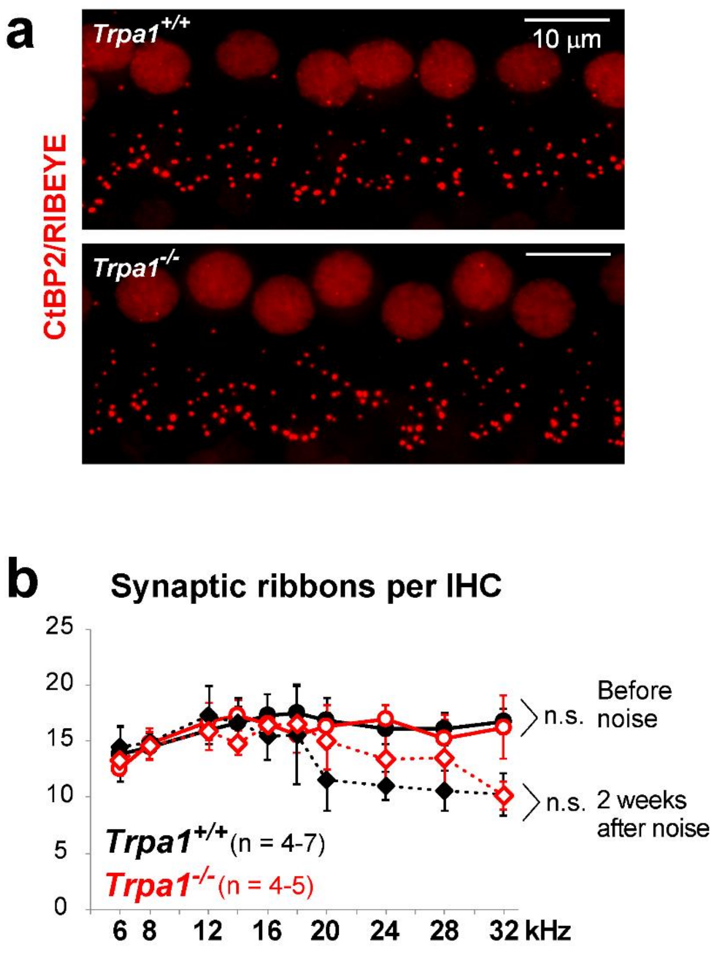


**Supplementary Figure 8.** ***TRPA1 deficiency does not exacerbate noise-induced loss of IHC ribbon synapses at high frequencies***

**(a)** Representative maximum-intensity projections of CtBP2/RIBEYE immunolabeling (red) in IHC from wild-type (top) and *Trpa1*$^{-/-}$ (bottom) mice at the 16 kHz cochlear region before noise exposure. **(b)** IHC synaptic ribbon counts along the cochlear length in wild-type (black) and *Trpa1*$^{-/-}$ (red) mice, before (circles, continuous lines) and two weeks after (diamonds, dotted lines) exposure to moderate 100 dB SPL broadband noise for 30 min. Data are shown as Mean ± SD (n.s. not significant, comparisons were made between grouped data by two-way ANOVA).
