## Supplementary material for "TRPA1 activation in non-sensory supporting cells contributes to regulation of cochlear sensitivity after acoustic trauma": Legends to Supplementary Videos

**Supplementary Video 1.** ***TRPA1-initiated long-lasting $Ca^{2+}$ responses in Hensen's cells of a wild-type mouse***

Time-lapse ratiometric imaging of $Ca^{2+}$ responses to the puff application of 200 µM of 4-HNE in a P7 wild-type cochlear explant (same as in Fig. 2a) shown at a 5X acceleration. The white arrow indicates the position, direction, and duration of the puff stimulation. Ratio images were calculated from fura-2 fluorescence at 340 and 380 nm excitation ($R = F_{340}/F_{380}$) and normalized to the baseline ratio value ($R_0$). $Ca^{2+}$ responses are shown using a pseudocolor scale that ranges between 1 (blue) and 2.5 (red) $R/R_0$. The size of the field of view is 104x104 µm. The original frame rate is 1.8 ratiometric pairs ($F_{340}/F_{380}$) per second. *IHC*, inner hair cells; *OHC*, outer hair cells; *HeC*, Hensen's cells; *CC*, Claudius' cells.

**Supplementary Video 2.** ***Propagation of TRPA1-initiated $Ca^{2+}$ responses from the Hensen's cells toward the Kolliker's organ***

Time-lapse video of the $Ca^{2+}$ responses to three consecutive puff applications of 400 µM of 4-HNE (from left to right) in a P1 wild-type cochlear explant shown at a 15X acceleration. The white arrows indicate the position, direction, and duration of each puff stimulation. All frames in the video represent the ratios of fura-2 fluorescence at 340 and 380 nm excitation ($R = F_{340}/F_{380}$) normalized to the baseline ratio value ($R_0$). $Ca^{2+}$ responses are shown using a pseudocolor scale for a range between 1 (blue) and 5 (red) $R/R_0$. The size of the field of view is 138x138 µm. The original frame rate is 2.2 ratiometric pairs ($F_{340}/F_{380}$) per second. *KoC*, Kolliker's organ cells; *IHC*, inner hair cells; *PC*, pillar cells; *OHC*, outer hair cells; *HeC*, Hensen's cells.

**Supplementary Video 3.** ***TRPA1 stimulation induced prominent tissue movements in the organ of Corti***

Bright field video recording at a 100X acceleration showing the tissue displacements induced upon stimulation with 200 µM 4-HNE in a wild-type cochlear explant (same as shown in Fig. 6a). The white arrows indicate the position, direction, and duration of the puff stimulation. The focal plane is located at the level of the OHC nuclei and pillar cell shafts. The video shows the original bright field imaging (*left*) as well as the subtracted frames that were 10 s apart (*right*). Pixels with an average gray value indicate no movement while darker and brighter pixels highlight the movement. The size of the field of view is 82x82 µm. *KoC*, Kolliker’s organ cells; *PC*, pillar cells; *OHC*, outer hair cells; *HeC*, Hensen’s cells. Age of the explant: P5.

**Supplementary Video 4.** ***Changes in the shape of outer pillar cells evoked by TRPA1 agonist***

Bright field video recording at a 40X acceleration showing the changes in the shape of outer pillar cells but not of OHCs upon stimulation with 200 µM 4-HNE in a wild-type cochlear explant (the same as shown in Fig. 6e). The white arrow indicates the position, direction and duration of the puff stimulation. The focal plane is located close to the pillar cell feet. The size of the field of view is 84x84 µm. *IPC*, inner pillar cells; *OPC*, outer pillar cells; *OHC*, outer hair cells. Age of the explant: P6.
